# Decoding Diets: Applying Non-Linear Machine Learning Models to Geometric Morphometric Analysis of Bovid Dental Mesowear Signatures

**DOI:** 10.64898/2026.04.14.718578

**Authors:** Rebecca Harbert, Kris Kovarovic, Ben Gruwier

**Affiliations:** Department of Anthropology, Durham University, Durham DH1 3LE, United Kingdom; Department of Physiology and Aging, University of Florida, Gainesville, FL, USA; Archaeology, Environmental Changes & Geo-Chemistry Research Group, Vrije Universiteit Brussel, Pleinlaan 2, 1050 Brussels, Belgium

## Abstract

Dental morphology and wear patterns provide insight into the dietary adaptations and ecological niches of living and extinct herbivores. Traditional classification statistics such as Linear Discriminant Analysis (LDA) are limited by assumptions of linearity, normality, and homoscedasticity. This study quantifies mesowear, the shape of molar cusps resulting from occlusal wear, and evaluates the performance of non-linear machine learning models in predicting herbivore diets based on geometric morphometric (GMM) data from adult mandibular second molars (M2) in bovids. We applied Generalized Procrustes Analysis and Principal Component Analysis (PCA) to digitized occlusal shape coordinates from 132 M2 specimens across 64 species. Using the resulting principal component scores, we compared the classification accuracy of LDA with three non-linear models: Random Forest, K-Nearest Neighbors, and Gradient Boosting. While LDA achieved a cross-validated accuracy of just 31%, all non-linear models achieved 99% cross-validation accuracy and 90% test accuracy, demonstrating substantially improved performance. Misclassification analyses revealed that non-linear models more effectively captured complex shape differences, particularly among species with overlapping wear patterns. Our findings support the integration of machine learning with geometric morphometrics to quantify mesowear and improve dietary classification, providing a framework for robust paleoecological inference.

## 1 Introduction

A central element in dietary reconstruction is the ability to infer diet from the morphology of molars, which for mammalian herbivores are both abundant and exceptionally well preserved in the fossil record (1–3). Teeth therefore provide one of the most direct proxies for understanding how both individual species and entire communities adapt dynamically to ecological pressures. Bovids, in particular, are especially informative due to their ecological diversity and rich fossil record, making them an important model for understanding dietary adaptations and environmental change (4,5). Situating these dietary signals within a broader environmental framework has become a major trend in paleontological research, linking events in mammalian evolution to ecological context (6–8).

Dietary behaviors are commonly inferred from occlusal dental wear, which can be evaluated on both microscopic and macroscopic scales. Microwear, assessed through two-dimensional scratches and pits or three-dimensional surface textures, captures short-term dietary signals (9,10), whilst qualitative and quantitative approaches to dental morphology yield insight into gross dietary adaptations. Distinct from these approaches, mesowear emphasizes cusp relief and occlusal morphology, reflecting the cumulative effects of attrition and abrasion over an individual’s lifetime (11). Mesowear analysis has proven particularly effective in distinguishing browsers, whose diets of leaves and softer vegetation maintain sharp, high-relief cusps, from grazers, whose consumption of abrasive grasses typically produces blunter and lower-relief occlusal surfaces (Figure 1) (11,12). The relative accessibility of mesowear, which does not require specialized equipment and integrates long-term dietary information, has made it a widely used method in paleoecological studies. However, mesowear remains a semi-quantitative and somewhat subjective approach, with inter-observer variability and reproducibility posing ongoing challenges (3). These limitations have encouraged the adoption of more quantitative and replicable frameworks such as geometric morphometrics (GM), which offers a rigorous alternative, providing a means to quantify shape variation while retaining the spatial relationships among landmarks (13–15). Traditional morphometrics, which reduce complex structures to distances, angles, or ratios, often oversimplifies shape and obscures spatial configurations (16). By analyzing landmark coordinates as Cartesian data, GM enables explicit encoding of both dimensional and spatial information, as well as direct visualization of shape differences (17).

**Fig 1.**
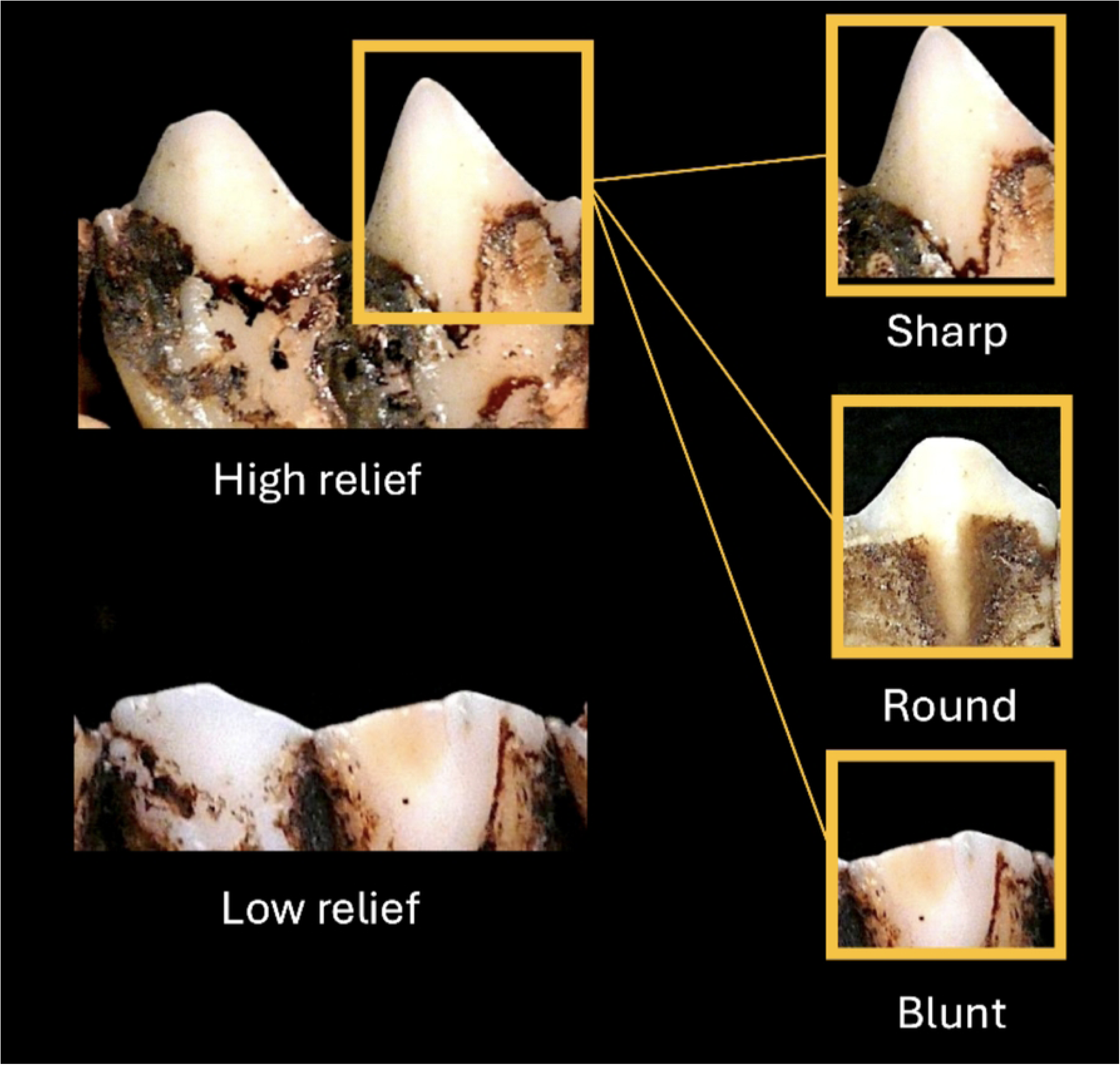
Examples of Cusp Relief and Shape on the Occlusal Surfaces of Molars in Browsing and Grazing Animals. Cusp relief and shape are key factors in mesowear scoring, as they reflect the degree of attrition and abrasion on the teeth. The image highlights differences in cusp height (high vs. low relief) and shape (sharp, rounded, or blunt cusps) typically observed in browsing and grazing animals. Browsers often exhibit higher, sharper cusps due to lower abrasion from softer vegetation, whereas grazers tend to have lower, more rounded or blunt cusps due to higher abrasion from abrasive grasses. Adapted from Mihlbachler et al., 2023, “Sharpening the Mesowear Score: Geometric Morphometric Analysis.”

Analyses of GM data commonly rely on multivariate techniques such as Principal Component Analysis (PCA) and Discriminant Function Analysis (DFA) (18). PCA is widely employed to reduce dimensionality and identify major axes of variation, facilitating both visualization and subsequent statistical analyses (19). DFA, particularly Linear Discriminant Analysis (LDA), has long been used as a classification tool in ecomorphological studies to distinguish functionally meaningful categories based on morphological variables (20). However, LDA depends on assumptions of multivariate normality and homogeneity of variance-covariance structures that are not always met in morphometric datasets (21). When these assumptions are violated, classification accuracy may be compromised, especially where dietary groups overlap morphologically. Thus, while LDA has played a central role in ecological and evolutionary studies, its reliance on parametric assumptions highlights the need for more flexible alternatives (22,23).

Machine learning (ML) provides such an alternative, offering approaches that are well suited to high-dimensional, non-linear data. Unlike traditional parametric models, ML algorithms do not rely on assumptions of normality or equal covariance, allowing them to capture more complex relationships within shape data (24,25). Random Forests, Gradient Boosting methods, and k-Nearest Neighbors are particularly relevant in this context, as they have demonstrated strong performance in biological and ecological applications (26,27). These models are robust to overfitting, accommodate large numbers of predictors, and can provide additional insights such as feature importance, helping to identify which aspects of morphology contribute most to classification outcomes (28,29).

Together, GM and ML create a promising framework for dietary reconstruction. GM supplies the quantitative description of morphological variation, while ML provides the analytical flexibility to classify categories without restrictive assumptions. By integrating these approaches, it becomes possible to refine predictions of herbivore diet and strengthen ecological interpretations based on fossil teeth. The present study applies this integrated framework to bovid mandibular second molars (M2), which display an intermediate degree of wear relative to M1 (more heavily worn) and M3 (least worn). This makes M2 informative for dietary comparisons, as it reflects consistent wear without the extremes observed in other positions (13). The entoconid cusp was chosen for analysis because of its stable wear patterns, comparative preservation, and availability across specimens.

We evaluate the performance of Random Forest, Gradient Boosting, and k-Nearest Neighbors against LDA in predicting dietary categories from GM-derived cusp morphology. By comparing classification accuracy and misclassification patterns across models, we assess the extent to which machine learning improves dietary prediction and explores the implications for paleoecological inference. In doing so, this study situates geometric morphometrics within a broader methodological toolkit that incorporates advanced computational approaches to the study of evolutionary and ecological morphology.

## 2 Methods

### 2.1 Bovid Sample and Data Collection

Our study analyzed 132 mandibular second molars (M2) representing 63 species of bovids from 12 natural history museum collections, with museum sources detailed in Supplementary Table 1; No permits were required for the described study, which complied with all relevant regulations. The dataset encompasses a broad taxonomic and ecological range, including members of the tribes Alcelaphini (n=4), Antilopini (n=20), Aepycerotini (n=4), Bovini (n=7), Caprini (n=6), Cephalophini (n=23), Hippotragini (n=6), Reduncini (n=5), Neotragini (n=20), Rupicaprini (n=9), Ovibovini (n=3), and Tragelaphini (n=18), along with minor representation from *N. batesi* and *R. rupicapra* groups. The number of individuals per species ranged from one to eight. Dietary assignments were based on Lintulaakso and Kovarovic (30) and include eight categories consolidated into four major ecological strategies—browsers (n=36), grazers (n=47), mixed feeders (n=37), and frugivores (n=12) (Supplementary Table 1; Supplementary Figure 1). These groupings provided a more balanced approach between ecological interpretability and statistical robustness, enabling meaningful comparison of shape signals across dietary strategies. Specimens were photographed in lingual view, with the occlusal surface oriented perpendicular to the camera lens. A 2 mm scale was included in all images. When right mandibular tooth rows were unavailable, left rows were mirrored to maximize sample size. This standardized photographic procedure ensured comparability of cusp outlines across individuals and species (Figure 2A).

**Fig 2.**
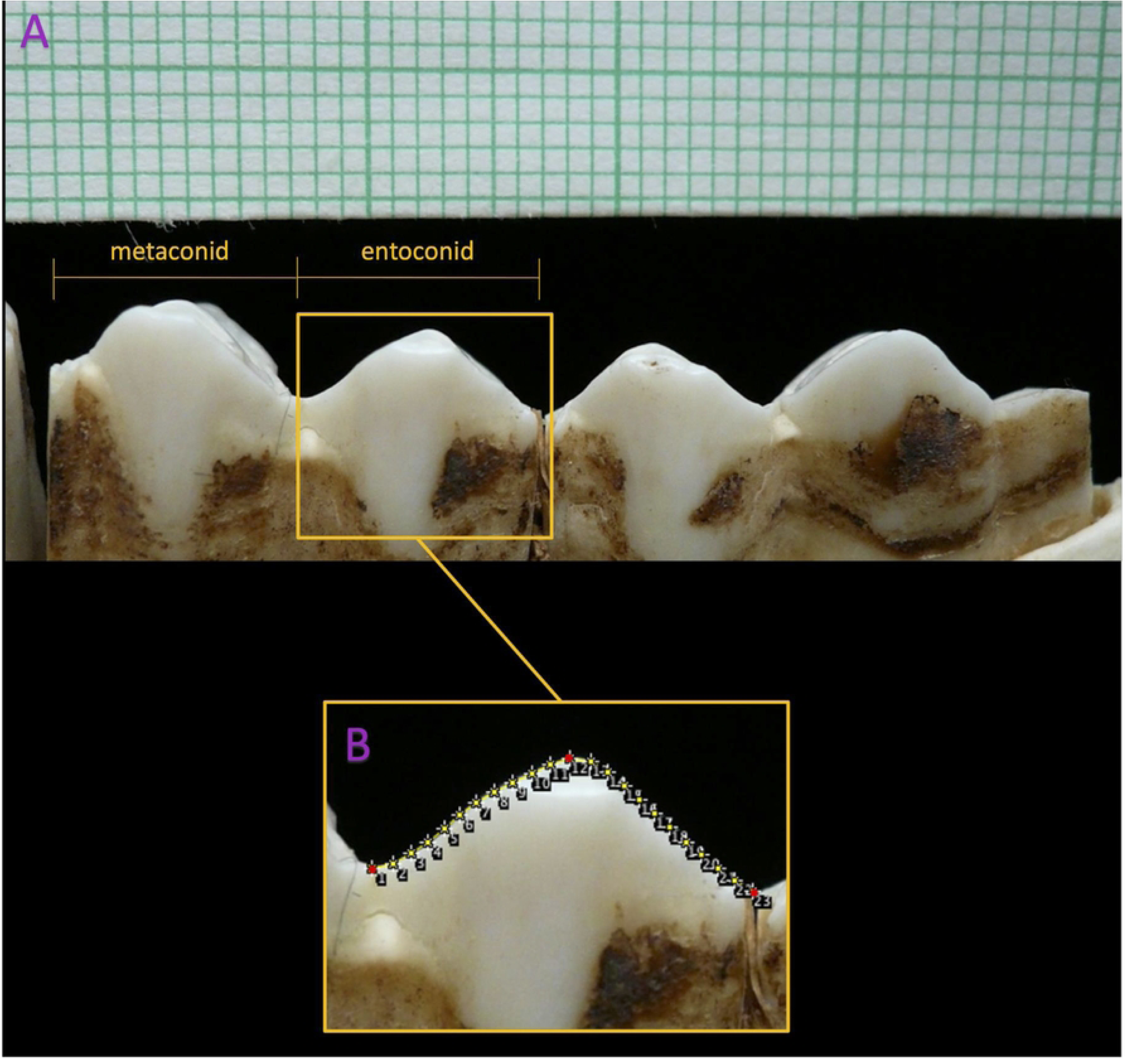
Lingual View of the Mandibular Second Molar (M2) of Damaliscus dorcas Showing Photographic Angle and Landmark Digitization. (A) Lingual view of the mandibular second molar (M2) of *Damaliscus dorcas* showing photographic angle. The specimen was oriented so the lingual wall of M2 was perpendicular to the camera lens and oriented where M2 is to the right of M3 and M3 is on the far left. (B) Digitization of the Occlusal Outline of the Entoconid Cusp on the Mandibular Second Molar (M2) of Damaliscus dorcas depicts the process of landmark digitization on the occlusal surface of the mandibular second molar (M2) on the same tooth row from *Damaliscus dorcas* as shown in Figure 2. Landmarks were placed along the outline of the entoconid cusp, beginning at the valley between the metaconid and entoconid cusps and continuing along the ridge to the distal edge of M2. The red points indicate the true anatomical landmarks: the first landmark at the valley between the cusps (landmark 1), the twelfth landmark marking the highest point on the entoconid (landmark 12), and the final landmark at the distal edge of the molar (landmark 23). The yellow points represent semi landmarks that were allowed to slide along the tangent to the curve to optimize alignment across specimens. The grid in the background provides a scale for measuring the outline.

### 2.2 Mesowear and Landmark Digitization

Mesowear analysis provides a semi-quantitative framework for dietary inference by transforming qualitative observations of cusp shape and relief into measurable categories (11,12,31). In its traditional implementation, mesowear scoring evaluates cusp shape (sharp versus rounded) and occlusal relief (high versus low), reflecting the balance between attritional and abrasive wear processes associated with different feeding strategies (11). Building on this foundation, the present study adopts the geometric morphometric approach of Mihlbachler et al. (13) to capture the same macroscopic wear features as continuous shape data. This landmark-based method provides a high-resolution, quantitative representation of mesowear, enabling finer evaluation of shape gradients associated with dietary strategy. The occlusal outline of the entoconid cusp was digitized for each specimen using ImageJ (32). Landmarks were placed beginning at the valley between the metaconid and entoconid, following the entoconid ridge, and terminating at the distal edge of M2 (Figure 2B). Following the protocol outlined by Mihlbachler et al. (13), a total of 23 landmarks were recorded: three fixed anatomical points (valley, cusp apex, and distal edge), and 20 sliding semilandmarks positioned along the cusp ridge (Figure 2B). Fixed landmarks correspond to biologically homologous points, while sliding semilandmarks capture curvature and were permitted to adjust position to minimize bending energy (15). Digitized coordinates (Supplementary Table 1) were first processed in Python to standardize file format and ensure compatibility with subsequent analyses. Generalized Procrustes Analysis (GPA) was then applied to remove variation related to translation, rotation, and scale, and Principal Component Analysis (PCA) was performed to summarize the main axes of shape variation. The resulting Procrustes-aligned shape coordinates and principal component (PC) scores were integrated with specimen metadata for downstream statistical modeling.

### 2.3 Shape Analysis

To remove variation due to orientation, position, and scale, landmark configurations were aligned using Generalized Procrustes Analysis (GPA), following established geometric morphometric procedures (33,34). Sliding semilandmarks were adjusted by minimizing thin-plate spline (TPS) bending energy (35), ensuring that placement reflected true morphological correspondence across specimens rather than arbitrary digitization. This procedure produced Procrustes coordinates representing the shape variation of each specimen in a common reference space. Principal Component Analysis (PCA) was applied to the Procrustes coordinates to reduce dimensionality and summarize axes of variation (19). The number of components retained for subsequent analyses was determined by inspection of the scree plot of eigenvalues, which indicated an inflection after the third principal component (see Figure 3). Because the aligned coordinates are already scaled and centered, no further normalization was required. Allometry, the influence of size on shape, can confound GM analyses (19). However, previous research demonstrates that the influence of size on cusp shape in bovids is minimal, yielding statistically insignificant results (13). Consequently, allometry was not considered a significant factor in this study.

**Fig 3.**
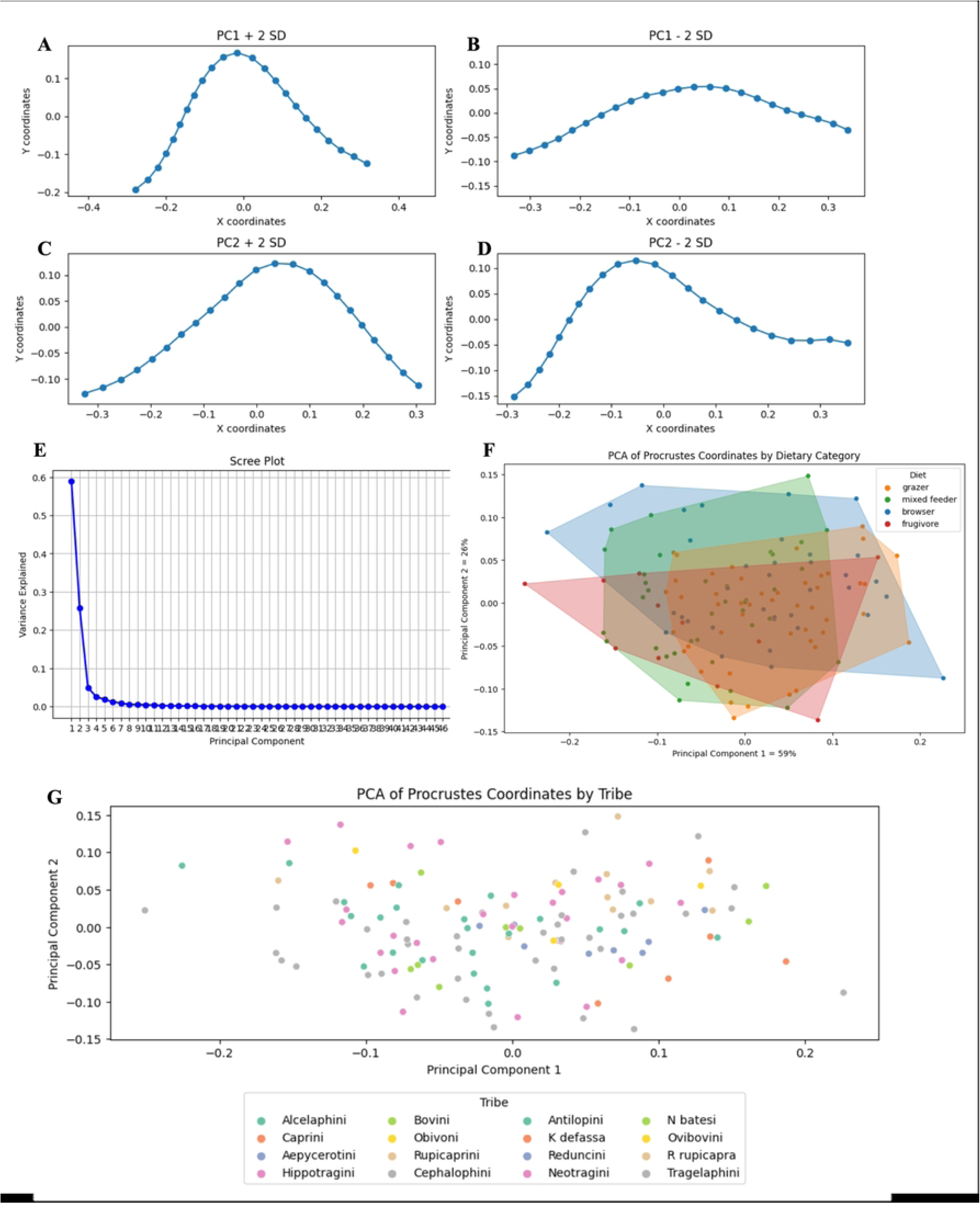
PCA of Procrustes Coordinates and Scree plot. (A-D) Shape Deformation Grids of Mandibular Second Molar (M2) Along PC1 and PC2. (A) PC1 + 2 SD shows cusp shapes at +2 standard deviations along PC1, where the cusps appear higher, indicating an increase in occlusal relief. (B) PC1 - 2 SD displays cusp shapes at −2 standard deviations along PC1, where the cusps are lower, suggesting a reduction in occlusal relief. The difference in elevation between these plots suggests that PC1 is related to changes in occlusal relief, with PC1 + 2 SD aligning with increased occlusal relief and PC1 - 2 SD with reduced occlusal relief. (C) PC2 + 2 SD shows shape changes at +2 standard deviations along PC2, possibly reflecting variations in the sharpness or positioning of the cusp apices. (D) PC2 - 2 SD depicts shape changes at −2 standard deviations along PC2, which may reflect the development of rounder and more blunt cusp shapes. (E) Scree plot (eigenvalue spectrum) used to determine the number of principal components retained. A clear elbow is observed at PCs 3–4, indicating a natural cutoff beyond which additional components contribute diminishing variance likely consistent with noise; PC1 explains 59% and PC2 explains 26% of total variance (85% cumulative) (F) PCA Plot of Procrustes Coordinates by Dietary Category. The x-axis represents the first principal component, which explains 59% of the total variance in the data, while the y-axis represents PC2, explaining 26% of the variance in the data. Each point on the scatter plot corresponds to an individual’s cusp shape, with colors indicating dietary category: orange for grazers, green for mixed feeders, blue for browsers, and red for frugivores. The addition of convex hulls around each dietary category provides a clear visual boundary for the data points for each group. (G) The PCA plot of Procrustes coordinates by tribe displays the distribution of mandibular second molar (M2) cusp shapes across various tribes as represented by the first two principal components. Each point is color-coded according to tribe, with certain tribes, such as Alcelaphini and Hippotragini, showing more distinct clustering patterns. Other tribes display a more dispersed distribution, with some overlap between groups. The principal components highlight the variation in cusp shapes, capturing differences both within and between the tribes.

### 2.4 Classification

Prior to classification, statistical assumptions were evaluated. All analyses were conducted using a between-subjects design, with each specimen treated as an independent observation. Normality of PCA scores was assessed using the Shapiro–Wilk test, which indicated significant deviations from normality. Log transformation of PCA scores was performed to improve normality; however, deviations persisted (Supplementary Figures 3 and 4). Homogeneity of covariance was tested with Box’s M, which was generally upheld (Supplementary Figure 5), while multicollinearity was reduced through PCA (Supplementary Figure 2). These checks underscored that Linear Discriminant Analysis (LDA) could be applied cautiously but also highlighted the potential benefits of non-parametric approaches less sensitive to these assumptions (23). Statistical significance was assessed at α = 0.05. No formal outlier removal was performed, and all observations were retained to preserve biological variability. The dataset contained no missing values after preprocessing; therefore, no imputation or exclusion procedures were required. As a traditional benchmark, Linear Discriminant Analysis (LDA) was implemented using PCA scores as predictors and dietary categories as response variables. LDA seeks linear combinations of features that maximize separation among groups, but its performance depends on adherence to parametric assumptions of multivariate normality and homogeneity of variance (21).

### 2.5 Machine Learning Models

To evaluate alternatives to LDA, three machine learning algorithms were employed: Random Forest, k-Nearest Neighbors, and Gradient Boosting. These models were chosen for their ability to model non-linear relationships and their established utility in ecological and morphometric applications (26,27). Random Forest is an ensemble method that constructs multiple decision trees from bootstrapped subsets of data and aggregates their predictions, reducing variance and improving generalizability (28). Gradient Boosting builds trees sequentially, each iteration correcting errors of the previous, allowing it to capture subtle, complex patterns (29). K-Nearest Neighbors classifies specimens based on the majority class of their closest neighbors in multivariate space, providing a simple yet effective non-parametric approach (36). The dataset was partitioned into training (80%) and testing (20%) subsets using stratified sampling to preserve class balance. The first three principal component scores, which together explained 90% of the total variance, served as input features. For machine learning models, hyperparameters were optimized using grid search with cross-validation. For example, Random Forest models were tuned for the number of trees and feature subsets, while Gradient Boosting was optimized for learning rate, tree depth, and minimum sample size per leaf. Model performance was assessed using accuracy, precision, recall, F1 score, and ROC AUC. Cross-validation provided estimates of model stability, while test set evaluation assessed generalization to unseen data. Learning curves were generated to evaluate performance trends as a function of training set size, allowing assessment of overfitting or underfitting. Misclassification analyses further examined species or dietary groups prone to error, with particular attention to overlaps between grazers and mixed feeders. Models were evaluated with stratified k-fold cross-validation and a held-out test set reported separately in Table 1. To avoid data leakage, all preprocessing (PCA, scaling) was fit within each training fold and applied to the held-out fold only. Hyperparameters for RF, k-NN, and GB were tuned by cross-validation using the training data only; the final model was refit on all training folds and evaluated once on the untouched test set. We additionally assessed learning curves to inspect the training–validation gap.

**Table 1.** Performance Metrics for Machine Learning Models. This figure compares the performance metrics of Random Forest, K-Nearest Neighbor (k-NN), and Gradient Boosting models. Each model shows high cross-validation mean accuracy, strong training and test accuracy, and high precision, recall, and F1 scores, with ROC AUC values ranging from 0.95 to 1.00, indicating robust and reliable classification performance.

| Model | Cross-Validated Mean Accuracy | Training Model Accuracy | Testing Model Accuracy | Precision | Recall | F1 | ROC-AUC |
| --- | --- | --- | --- | --- | --- | --- | --- |
| Random Forest | 0.99 ±0.02 | 1.00 | 0.90 | 0.90 | 0.90 | 0.90 | 0.97 |
| K-Nearest Neighbor | 0.99 ±0.02 | 0.99 | 0.90 | 0.90 | 0.90 | 0.90 | 0.95 |
| Gradient Boost | 0.99 ±0.02 | 1.00 | 0.90 | 0.90 | 0.90 | 0.90 | 1.00 |
| LDA | 0.37 ±0.09 | 0.38 | 0.37 | 0.23 | 0.37 | 0.25 | 0.67 |

## 3 Results

### 3.1 Shape Analysis and Linear Discriminant Analysis

The PCA of Procrustes-aligned mandibular second molar (M2) coordinates revealed major axes of shape variation consistent with dietary differentiation (Figure 3A-D). The first three principal components together explained approximately 90% of the total variance, with PC1 accounting for 59%, PC2 for 26%, and PC3 for an additional 5%. PC1 reflected variation in cusp relief and enamel ridge development, while PC2 captured relative basin depth and hypoconulid extension (Figure 3A–E). The scree plot indicated a sharp inflection after the third component, suggesting these axes provide the most informative summary of shape variation (Figure 3E). When plotted by dietary category, browsers and grazers showed partial separation along PC1, with mixed-feeders occupying intermediate positions (Figure 3F). Examination of shape variation by tribe revealed phylogenetic clustering, underscoring the influence of lineage on molar morphology (Figure 3G). The PCA-by-tribe ordination further shows that tribal affiliation provides a secondary axis of structure in the data, with specialized tribes (e.g., Alcelaphini grazers, Tragelaphini browsers) exhibiting clearer morphological cohesion than generalist tribes (e.g., Bovini, Reduncini).

LDA was applied to the PCA-transformed shape data, using the first three principal component (PC) scores as input variables to assess classification accuracy of dietary groups. The model achieved moderate separation of browsers and grazers, with mixed-feeders distributed between them. Cross-validation accuracy was 0.31 ± 0.14, with training and test accuracies of 0.38 and 0.37, respectively. Model performance metrics included a precision of 0.23, recall of 0.37, F1 score of 0.25, and ROC AUC of 0.68. Misclassifications primarily occurred between mixed-feeders and the browsing and grazing categories (Figure 4A; Table 1). The first discriminant axis reflected cusp height and basin constriction, while the second emphasized enamel ridge curvature. Results are summarized in the LDA scatterplot and accompanying confusion matrix (Figure 4; Table 2). Analysis of misclassified specimens provided insight into morphological overlap among dietary groups. Most errors involved mixed-feeders, whose molar shapes overlapped extensively with both browsers and grazers (Supplementary Figure 6). In several cases, specimens classified as browsers exhibited grazing-like hypoconulid reduction, while some misclassified grazers retained basin depth resembling that of browsers.

**Fig 4.**
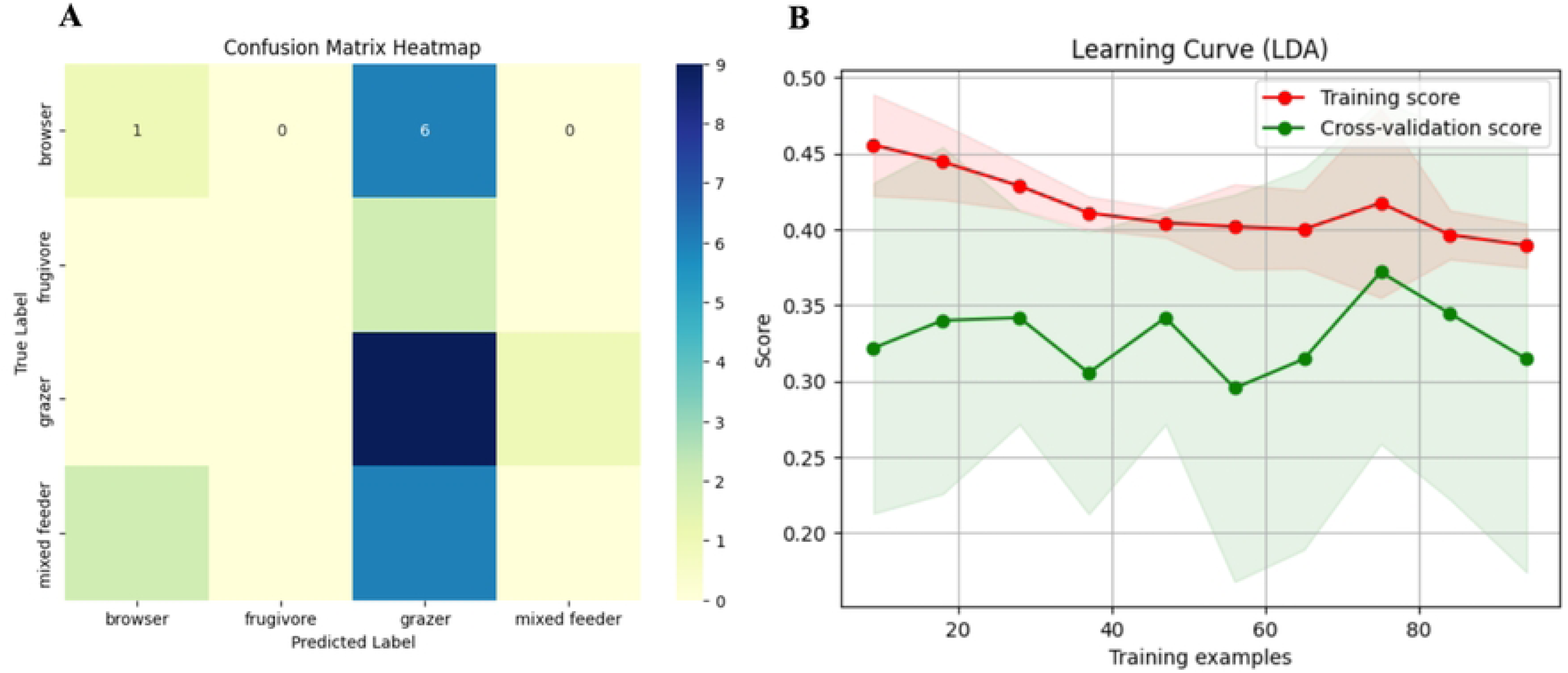
Performance Metrics for Linear Discriminant Analysis. (A) Confusion Matrix Heat Map for LDA. The heatmap visualizes the confusion matrix, showing the frequency of correct (diagonal) and incorrect (off diagonal) predictions. The LDA model exhibits significant misclassification across all four dietary categories. (B) LDA Learning Curve. The above learning curve gives insights into the model’s performance on the training data (red line) versus the cross-validation data (green line). The significant drop in the cross-validation score relative to the training score suggests overfitting, where the model performs well on the training data but poorly on unseen data. The training examples along the x axis indicate the epoch count for cross-validation.

**Table 2.** Condensed misclassification table across classifiers (LDA, RF, k-NN, GB). Columns list species, tribe, true dietary label, misassigned label(s), and the model(s) responsible. Errors are chiefly non-grazers predicted to be grazers, reflecting overlap in PC-derived shape features. Consistently misclassified taxa (appearing in ≥2 models) include: *Aepyceros melampus* (Aepycerotini), *Bison bison* (Bovini), *Hippotragus equinus* (Hippotragini), and *Procapra pictaudata* (Antilopini). Abbreviations: LDA, Linear Discriminant Analysis; RF, Random Forest; k-NN, k-Nearest Neighbors; GB, Gradient Boosting

| Misclassification Table by Species |  |  |  |  |  |
| --- | --- | --- | --- | --- | --- |
| Species | Tribe | True label | Predicted labels | Models that misclassified | N errors |
| *Aepyceros*<br>*melampus* | Aepycerotini | Mixed-feeder | Browser,<br>Grazer | LDA,<br>Random<br>forest, K-Nearest<br>Neighbor,<br>Gradient<br>Boosting | 4 |
| *Bison bison* | Bovini | Grazer | Mixed-feeder | LDA,<br>Random<br>forest, K-Nearest<br>Neighbor,<br>Gradient<br>Boosting | 4 |
| *Cephalophus*<br>*leucogaser* | Cephalophini | Mixed-feeder | Grazer | LDA | 2 |
| *Cephalophus*<br>*natalensis* | Cephalophini | Frugivore | Grazer | LDA | 1 |
| *Hippotragus equinus* | Hippotragini | Mixed-feeder | Browser, Grazer | LDA, Random forest, K-Nearest Neighbor, Gradient Boosting | 4 |
| *Nesotragus moschatus* | Neotragini | Browser | Grazer | LDA | 1 |
| *Oryx gazella* | Hippotragini | Mixed-feeder | Grazer | LDA | 1 |
| *Ovibus moschatus* | Ovibovini | Mixed-feeder | Grazer | LDA | 1 |
| *Ovis dalli* | Caprini | Mixed-feeder | Grazer | LDA | 1 |
| *Philantomba monticola* | Cephalophini | Frugivore | Grazer | LDA | 1 |
| *Procapra picticaudata* | Antilopini | Browser | Grazer,<br>Mixed-feeder | LDA,<br>Random forest, K-Nearest Neighbor, Gradient Boosting | 4 |
| *Raphicerus campestris* | Neotragini | Browser | Grazer | LDA | 1 |
| *Raphicerus sharpie* | Neotragini | Browser | Grazer | LDA | 1 |
| *Sylvicapra grimmia* | Cephalophini | Mixed-feeder | Grazer | LDA | 1 |
| *Tragelaphus eurycerus* | Tragelaphini | Browser | Grazer | LDA | 1 |
| *Tragelaphus scriptus* | Tragelaphini | Browser | Grazer | LDA | 1 |

### 3.2 Machine Learning Classifiers

To evaluate non-linear approaches, Random Forest, k-Nearest Neighbors, and Gradient Boosting were trained on the same PCA-transformed data, using the first three principal component scores as input variables. Across models, cross-validated mean accuracy was 0.99– 1.00 with test accuracy around 0.90 (Table 1). Precision, recall, and F1 each ranged 0.95–1.00, with ROC-AUC 0.95–1.00. For the tree-based models (RF, GB), permutation-based importance identified PC1 and PC2 as consistently influential; model-agnostic permutation for k-NN yielded a similar ranking. The learning curves show rapid convergence with a small training–validation gap (Figure 5). Taken together with the stratified CV + held-out test design and leakage-free preprocessing, this argues against overfitting even with near-ceiling CV accuracy (Table 1; Figure 5). These results indicate that non-linear classifiers outperform LDA in capturing dietary signal.

**Fig 5.**
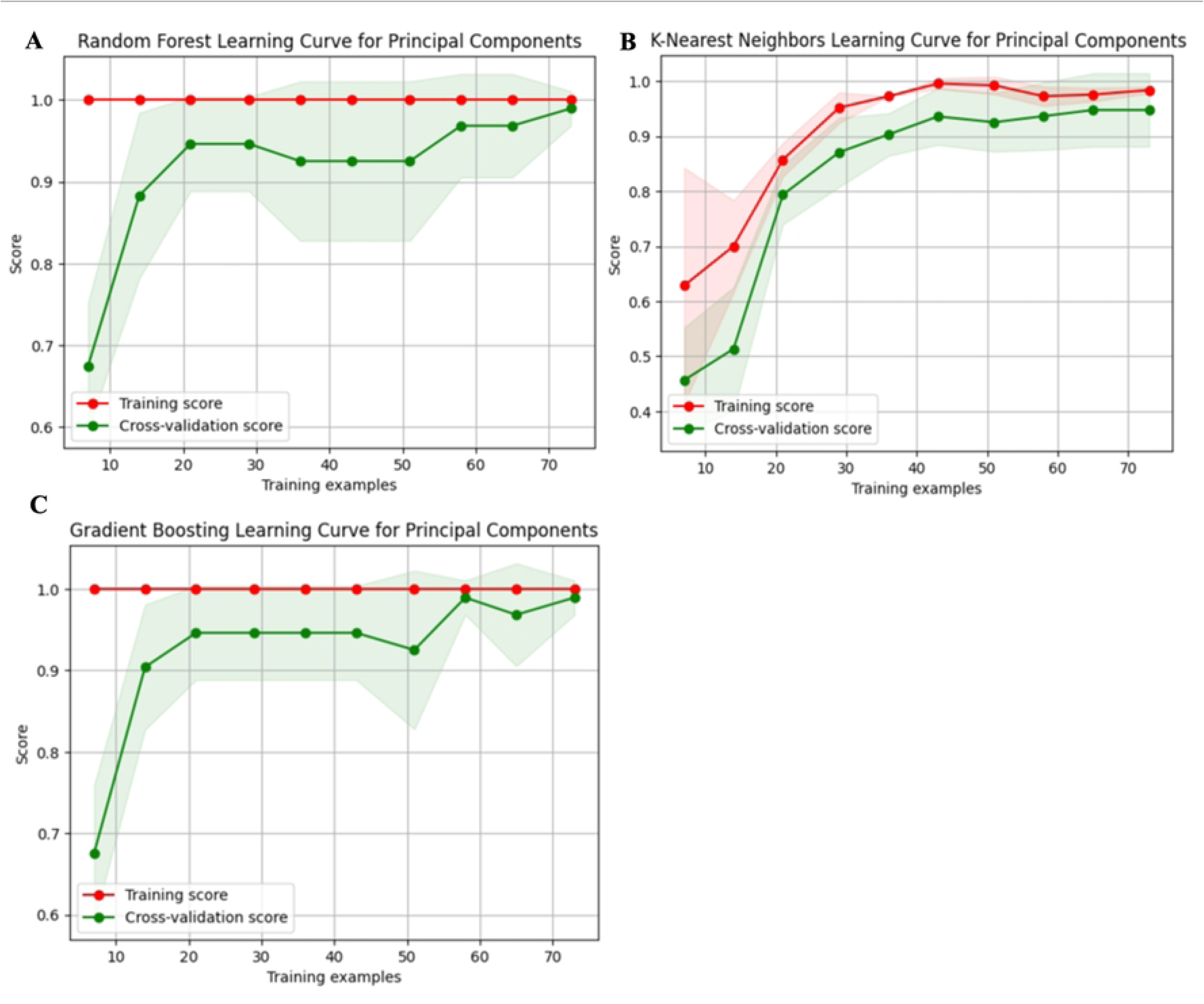
Learning curves for machine-learning classifiers trained on principal component scores of shape data. (A) Random Forest, (B) k-Nearest Neighbors, (C) Gradient Boosting. Red = training score; green = cross-validation (CV) score; shaded bands = ±1 SD across folds. (A) Random Forest: the training score remains consistently high; the CV score increases with more training examples and then stabilizes; as sample size grows, the CV curve converges toward the training curve, indicating reduced variance and improved model stability. (B) k-Nearest Neighbors: the training score shows a steep early rise and stabilizes near perfect accuracy; the CV score improves substantially with additional examples and eventually converges with the training score; both curves stabilize as the number of training examples increases. (C) Gradient Boosting: the training score remains consistently high; the CV score rises and then stabilizes; with more training data, the CV curve approaches the training curve, demonstrating improved and stable predictive performance. (x-axis: number of training examples; y-axis: score.)

Misclassifications for the non-linear models showed a complementary pattern: Random Forest and Gradient Boost most frequently labeled mixed-feeders as grazers, whereas k-Nearest Neighbors more often produced mixed-feeders as browsers, with occasional browser-to-grazer and grazer-to–mixed-feeder errors across models. These results reinforce that intermediate morphologies are the principal source of error (Table 2). Together with the overall gains in predictive accuracy, this indicates that non-linear classifiers capture the dietary signal more effectively than LDA while retaining similar error structure concentrated in mixed-feeder specimens.

### 3.3 Model Comparison Summary

A comparison of model performance underscores the trade-offs between linear and non-linear classifiers. LDA provided interpretable axes of shape differentiation but was limited by overlap in intermediate morphologies. Random Forest and Gradient Boosting achieved higher predictive accuracy, with Random Forest showing the best overall balance of interpretability and performance. K-NN underperformed relative to other methods. (Table 2; Figure 5).

## 4 Discussion

### 4.1 Model Performance and Methodological Considerations

The comparative evaluation of traditional and machine learning methods for dietary classification based on geometric morphometric (GM) data reveals insights into the strengths, limitations, and underlying mechanisms that contribute to their performance. By employing Principal Component Analysis (PCA), Linear Discriminant Analysis (LDA), and machine learning models including Random Forest, K-Nearest Neighbors (k-NN), and Gradient Boosting, this study provides a comprehensive assessment of the efficacy of these approaches in handling the complex shape data derived from the mandibular second molar of bovids.

In this study, PCA captured most of the total shape variance by the first three principal components, which effectively summarized the dominant axes of shape variation. The use of PCA facilitated dimensionality reduction by allowing high-dimensional landmark data to be represented by a small number of orthogonal variables that retain the majority of biologically meaningful shape information (Mitteroecker and Gunz, 2009). This reduction in dimensionality not only facilitated visualization but also made the data more manageable for subsequent analyses, such as LDA and machine learning models (Neha, 2015; Raschka, 2015). However, it is important to recognize that PCA outputs are statistical constructs derived from the data and do not necessarily correspond to direct biological processes (Mitteroecker & Gunz, 2009). The principal components represent directions of greatest variance, which may or may not align with biologically meaningful differences. Thus, while PCA remains an effective tool for data reduction and visualization, caution is warranted when interpreting results, as some variation may reflect noise or non-biological structure (Mitteroecker & Gunz, 2009). Additionally, the linear nature of PCA, while a strength in terms of interpretability and computational efficiency, can also be a limitation. Since PCA only captures linear relationships, it may not fully account for non-linear interactions within the data, potentially missing subtle but important variations in shape (Hastie et al., 2009). Nevertheless, PCA remains a crucial step in preprocessing GM data, as it helps to mitigate issues related to multicollinearity and improves the stability of downstream models by focusing on the most informative features (Mitteroecker and Gunz, 2009).

Linear Discriminant Analysis was applied as a traditional parametric approach to classify dietary groups based on the PCA-transformed coordinates. Although LDA identifies linear combinations of variables that maximize separation among groups (James et al., 2013; Kovarovic et al., 2011), its performance in this study was limited, achieving a mean cross-validation accuracy of only 31%. This underperformance likely reflects LDA’s reliance on assumptions of normality and homoscedasticity and its reduced capacity to capture non-linear relationships inherent in geometric morphometric data, since these conditions were not fully met in the current study (Kovarovic et al., 2011). The complexity and non-linearity of the shape data likely led to poor separation between dietary groups, as LDA’s linear discriminants could not adequately capture the true structure of the data. Furthermore, despite weighing each class, LDA’s sensitivity to imbalanced datasets and outliers may have exacerbated its struggles in this study, as even small deviations from the ideal conditions can significantly impact the model’s accuracy (Adams et al., 2004).

In contrast, machine Learning models demonstrated a markedly superior ability to classify dietary groups, offering a more flexible and robust approach compared to LDA. These models, including Random Forest, K-Nearest Neighbors (k-NN), and Gradient Boosting, leverage non-linear decision boundaries and ensemble learning techniques, which allow them to capture complex patterns in the data that LDA cannot (Hastie et al., 2009). Random Forest performed exceptionally well in classifying dietary groups, achieving a cross-validation accuracy of 99%. The model effectively managed the complexity of the shape data, capturing both linear and non-linear relationships without requiring strict parametric assumptions. This flexibility allowed Random Forest to identify subtle shape variation that LDA’s linear boundaries failed to capture. Feature importance analysis indicated that PC1 and PC2 contributed most to classification performance, suggesting that the primary axes of shape variation contained the strongest dietary signal. Importantly, the ability to extract feature importance values also provides interpretability, allowing identification of the specific shape components most relevant to dietary differentiation. Moreover, the absence of a widening gap between training and validation curves, together with consistent performance on the held-out test set, indicates that model accuracy was not the result of overfitting but reflected genuine generalization to unseen data. These findings underscore Random Forest’s capacity to model complex patterns in cusp shape while maintaining a high level of predictive accuracy.

The k-NN model, a non-parametric method that classifies a data point based on the majority class of its nearest neighbors, also performed exceptionally well, achieving a cross-validation accuracy of 99% (Altman, 1992). Its strength lies in capturing local structure within the data, allowing specimens with similar shapes to be grouped effectively without relying on predefined model structure. In this study, careful tuning of the number of neighbors and distance metric was necessary to optimize model accuracy. Because the model relies on distance-based comparisons, it effectively captured local patterns within the PCA-reduced shape data, grouping specimens with similar cusp shapes and dietary affiliations. This property was particularly useful given that the dataset exhibited partial clustering by diet and tribe, allowing k-NN to perform well even without linear decision boundaries. However, as with most distance-based algorithms, k-NN remains sensitive to the “curse of dimensionality,” where distances lose discriminative power as feature space expands (Hastie et al., 2009). PCA helped mitigate this issue by reducing the data to a lower-dimensional, biologically meaningful representation that retained the principal axes of shape variation (Mitteroecker & Gunz, 2009; Zelditch et al., 2012). This balance between dimensionality reduction and local sensitivity likely contributed to the model’s strong classification performance. Misclassifications were primarily observed in mixed feeders, reflecting true overlap in cusp shape rather than model error.

Gradient Boosting, another ensemble method, sequentially builds models to correct the errors made by previous ones, thereby reducing bias and variance (Friedman, 2001). The Gradient Boosting model achieved a cross-validation accuracy of 99% and demonstrated strong generalization capabilities. Regularization parameters such as learning rate and tree depth were tuned through cross-validation to limit overfitting. The convergence of training and validation accuracy (Figure 5) and the consistency between cross-validation and test performance suggest that the model generalized appropriately to unseen data. The iterative nature of Gradient Boosting, where each new model focuses on the hardest-to-predict instances, allows it to capture complex, non-linear relationships within the data that other models might miss (Friedman, 2001). The strength of Gradient Boosting lies in its ability to create a highly accurate model by combining the strengths of many weak learners (typically decision trees) (Friedman, 2001). By minimizing the loss function at each step, Gradient Boosting fine-tunes the model to better predict the outcomes, leading to superior performance in challenging datasets, such as the PCA-reduced cusp shape data analyzed in our study (Friedman, 2001).

Taken together, the contrast between the performance of LDA and the machine learning models underscores the limitations of traditional parametric methods in the context of geometric morphometrics. Linear decision boundaries and strict distributional assumptions limit its effectiveness in capturing the complex, non-linear relationships present in GM data (James et al., 2013; Zelditch et al., 2012). In contrast, the machine learning models in this study achieved markedly higher accuracy and captured relationships between cusp shape and diet that LDA’s linear approach could not resolve. The ensemble nature of Random Forest and Gradient Boosting, combined with the simplicity and adaptability of k-NN, allowed these models to achieve near-perfect accuracy, demonstrating their superior ability to handle the intricacies of morphological data.

Despite this strong performance, it is important to acknowledge that machine learning models introduce practical considerations related to model complexity, computational cost, and the potential for overfitting, particularly when applied to morphometric datasets of moderate size (Hastie et al., 2009). While careful tuning and validation are essential for minimizing overfitting and ensuring reliable performance, differences among models reflect inherent trade-offs between predictive flexibility, interpretability, and computational demands. Among the models tested, Random Forest provided the most consistent and interpretable performance, while Gradient Boosting achieved similar accuracy at greater computational cost. The k-NN model also performed well but showed slightly higher sensitivity to parameter selection, highlighting trade-offs between simplicity and stability across approaches.

### 4.2 Ecological Implications

The performance of the classification models provides important observations about the wear-related variation in bovid molar shape. The machine learning models demonstrated a high level of accuracy in classifying dietary categories based on the shape of the mandibular second molar. These models’ ability to accurately distinguish between different dietary groups confirms that specific dental shapes are strongly indicative of particular diets (Fortelius & Solounias, 2000; Kaiser, 2003; Semprebon et al., 2019; Mihlbachler et al., 2023). The high accuracy observed in the classification of browsers supports the interpretation that these species possess distinct shape traits that are effectively captured by the models. The sharper and higher-relief cusps associated with browsers, as indicated by the correct classifications, were key features identified by the models as characteristic of a browsing diet, confirming patterns reported in previous mesowear studies (Ackermans, 2020; Fortelius & Solounias, 2000). This observation aligns with the understanding that these dental traits are well-suited for processing soft, fibrous vegetation (Fortelius and Solounias, 2000; Schubert et al., 2006). Similarly, each machine learning models’ ability to correctly classify grazers, which was reflected in their performance metrics (Figure 5; Tables 1 and 2), suggests that lower, more rounded cusps are reliably indicative of a diet primarily composed of abrasive grasses. The success of the models in distinguishing grazers from other dietary groups implies that these shape features are sufficiently distinct to be recognized, even in a complex dataset. This observation supports the idea that the evolution of these dental traits in grazers is an adaptation to the mechanical demands of grinding tough, fibrous plant material (Fortelius and Solounias, 2000; Semprebon et al., 2019).

The overlap in classification performance observed for mixed feeders, where some misclassifications occurred, probably reflects a more generalized dental shape of these species. The models’ occasional difficulty in distinguishing mixed feeders from other dietary categories suggests that these species possess intermediate cusp shapes, which could be indicative of a flexible diet that includes both browse and graze. This observation highlights the flexibility of mixed feeders in response to varying ecological conditions, which is likely also reflected in their less specialized dental shapes (Damuth and Janis, 2011; T. A. Franz-Odendaal and Solounias, 2004). These misclassification trends suggest that intermediate shapes blur categorical boundaries, consistent with flexibility in feeding strategies (Table 2). Furthermore, misclassification patterns differed among models and were most pronounced between browsers and mixed feeders or grazers (Table 2). In particular, some browsers were consistently assigned to mixed-feeding or grazing categories, suggesting overlap in dental wear patterns among these dietary groups. Although these categories remain distinct, such overlap may reflect similar functional responses to comparable dietary or environmental conditions (Rivals et al., 2010).

The overall performance of the classification models also underscores the potential for using advanced machine learning techniques to enhance our understanding of the ecological roles of extinct species. The ability of these models to accurately classify dietary groups based on dental shape suggests that similar approaches could be applied to fossil data, providing more precise reconstructions of past environments and dietary behaviors (Kovarovic et al., 2011; Zelditch et al., 2012).

### 4.3 Future Directions and Limitations

The findings of this study highlight several promising directions for expanding the current framework. A logical next step would be to extend the shape coordinate analyses to include additional molars, such as the first and third mandibular molars, or to incorporate full occlusal outlines of the tooth row to capture a more comprehensive view of wear. It would also be valuable to cross-reference these results with other dietary proxies, such as dental microwear and isotopic data, to test the consistency of shape-based classifications across independent lines of evidence. In addition, the application of image-based deep learning approaches, such as convolutional neural networks, may offer opportunities to automate feature extraction and improve the scalability of shape-based dietary reconstructions.

At the same time, several limitations should be acknowledged when interpreting the high classification accuracies observed in this study. Although training, validation, and test performance were closely aligned across models, suggesting limited overfitting, the near-ceiling accuracy achieved by the machine learning approaches likely reflects both a strong dietary signal in mandibular second molar shape and characteristics of the dataset itself, including sample size, class structure, and partial clustering by diet and tribe. As a result, classification performance may be optimistically estimated within this dataset, and model generalizability should be evaluated through application to independent or fossil datasets. Future work incorporating larger and more taxonomically diverse samples will be critical for assessing the robustness of these models and for distinguishing true biological signal from dataset-specific structure.

## 5 Conclusions

Machine learning models were applied to the geometric morphometric analysis of mandibular second molar (M2) cusp outlines in bovid species, with the goal of classifying these species into dietary categories based on their mesowear signal. The analysis demonstrated that models such as Random Forest, Gradient Boosting, and K-Nearest Neighbors outperformed traditional Linear Discriminant Analysis (LDA) in capturing the complex non-linear relationships present in the shape data. These models generally succeeded in distinguishing between grazers and browsers, suggesting that geometric morphometric data of cusp outlines can provide meaningful insights into dietary adaptations.

While the findings are promising, they should be interpreted cautiously. The ability of these models to distinguish dietary categories confirms previously demonstrated correlations between cusp shape and diet, though further research is necessary to strengthen these connections. The results indicate that machine learning approaches have the potential to improve on traditional methods in dietary reconstruction, particularly in cases where subtle morphological differences may not be easily captured through conventional techniques.

This research contributes to the growing body of knowledge on the application of geometric morphometrics and machine learning in paleoecology, illustrating how these tools can be used to explore evolutionary and ecological dynamics in herbivorous mammals. However, the need for continued refinement of the models to improve their accuracy and reliability is evident. Future work should focus on enhancing these models, possibly by incorporating additional morphological or ecological data, to fully realize their potential in reconstructing the diets of both extant and extinct species.

In conclusion, while valuable insights have been gained into the use of machine learning models in dietary classification, there is also a clear emphasis on the need for further validation and refinement. The integration of geometric morphometrics with advanced machine learning techniques offers a promising avenue for future research, with the potential to deepen our understanding of the evolutionary history and ecological roles of herbivorous mammals.

## Data Code and Availability

The XY coordinates used for data processing and modeling are available in Supplementary Table 1. The complete source code used for data processing, modeling, and analysis is available in the following GitHub repository: https://github.com/Harbert0/gm-m2-morphometrics-pipeline. Original images are available upon request.

## Acknowledgments

Thanks go to many people who facilitated data collection on extant bovid dentition in Europe and the US: The Natural History Museum, London: Paula Jenkins, Daphne Hills, Louise Tomsett, Rob Kruszynski; Smithsonian National Museum of Natural History, Washington DC: Linda Gordon; American Museum of Natural History, NYC: Bob Randall, Eileen Westwig, Neil Duncan; Field Museum, Chicago: Bill Stanley; Powell-Cotton Museum, Kent, UK: Malcolm Harmon; Zoological Museum, Copenhagen, Denmark: Hans Baagoe, Mogens Andersen; Swedish Museum of Natural History, Stockholm: Per Ericson, Olavi Gronwall, Lars Werdelin; Natural History Museum Vienna, Austria: Barbara Herzig, Helen Jousse, Alexander Bibl; Naturalis, Leiden, The Netherlands: Lars van der Hoek Ostende, John de Vos, Hein van Grouw; Museum of Natural History, Berlin, Germany: Frieder Meyer, Detlef Willborn, Irene Mann; Royal Museum of Central Africa, Tervuren, Belgium: Emmanuel Gilissen, Wim Wendelin, Garin Cael; Hungarian Natural History Museum, Budapest: Gabor Csorba, Laszlo Peregovits, Zoltan Vos. Funding to KK for data collection was provided by The Leverhulme Trust and the SYNTHESYS Project (http://www.synthesys.info/), which was financed by European Community Research Infrastructure Action under the FP6 “Structuring the European Research Area” Programme and the Royal Society International Joint Project. BG wishes to acknowledge FWO for a postdoctoral grant (1275824N).

## Supporting Information

**S1 Fig. Class Distribution Histogram.** The histogram displays the number of instances for each dietary category in the dataset. The Y axis represents the count or the number of individuals who are classified in a particular category. The number at the top of each bar represents the number of individuals for the respective dietary category with grazers (orange) containing 47 individuals, mixed feeders (green) containing 37 individuals, browsers (blue) containing 36 individuals, and frugivore (red) containing 12 individuals. Browsers are slightly overrepresented, while frugivores are underrepresented. This distribution informed the weighting adjustments applied during the DFA to maintain analytical rigor.

**S2 Fig. Correlation Matrices for Procrustes Coordinates (A) and Principal Components (B).** S2.A illustrates **a** heatmap showing the high level of multicollinearity among the original shape coordinates, with many pairs of coordinates showing strong correlations. S2.B shows a correlation matrix of the principal components. After PCA transformation, the multicollinearity is removed, as indicated by the near-zero correlations between the principal components, making the PCA transformed coordinates a suitable candidate for use in DFA.

**S3 Fig. Histogram of p-values for Shapiro-Wilk Test (Original PCs).** The histogram shows that most p-values are near zero, indicating a significant violation of the normality assumption for the original principal components

**S4 Fig. Histogram of p-values for Shapiro-Wilk Test (Log-transformed PCs).** Even after log transformation, the p-values remain skewed towards zero, suggesting that non-parametric methods may be better suited for analysis.

**S5 Fig. Covariance Matrices by Class.** The heatmaps show the covariance matrices for each dietary class, indicating low variance and covariance, which suggests that LDA is more appropriate than QDA for this dataset. S5.A shows the heatmap for browsers, S5.B for mixed feeders, S5.C for frugivores, and S5.D for grazers.

**S6.Fig. Original Images of the 4 most misclassified samples within the LDA, Random Forest, k-NN, and Gradient Boosting models.** S6.A shows the tooth row for *Hippotragus equinus,* S6.B shows the tooth row for *Procapra picticaudata,* S6.C shows the tooth row for *Aepyceros melampus,* and S6.D shows the tooth row for *Bison bison*.

